# Deciphering copper coordination in the animal prion protein amyloidogenic domain

**DOI:** 10.1101/507624

**Authors:** Giulia Salzano, Martha Brennich, Giordano Mancini, Thanh Hoa Tran, Giuseppe Legname, Paola D’ Angelo, Gabriele Giachin

## Abstract

Prions are pathological isoforms of the cellular prion protein (PrP^C^) responsible for transmissible spongiform encephalopathies (TSE). PrP^C^ interacts with copper through unique octarepeat and non-octarepeat (non-OR) binding sites. Previous works on human PrP^C^ suggest that copper binding to the non-OR region may have a role during prion conversion. The molecular details of copper coordination within the non-OR region are not well characterized. By means of small angle X-ray scattering (SAXS) and extended X-ray absorption fine structure (EXAFS) spectroscopy, we have investigated the Cu(II) structural effects on the protein folding and its coordination geometries when bound to the non-OR region of recombinant PrP^C^ (recPrP) from animal species considered high or less resistant to TSE. As TSE-resistant model, we used ovine PrP^C^ carrying the protective polymorphism at residues A136, R154 and R171 (OvPrP ARR); while as highly TSE-susceptible PrP^C^ models we employed OvPrP with polymorphism V136, R154 and Q171 (OvPrP VRQ) and Bank vole recPrP (BvPrP). Our results reveal that Cu(II) affects the structural plasticity of the non-OR region leading to a more compacted conformation of recPrP. We also identified two Cu(II) coordinations in the non-OR region of these animal species. In *type-1* coordination present in OvPrP ARR, Cu(II) is coordinated by four residues (S95, Q98, M109 and H111). Conversely, the *type-2* coordination is present in OvPrP VRQ and BvPrP, where Cu(II) is coordinated by three residues (Q98, M109 and H111) and by one water molecule, making the non-OR region more flexible and open to the solvent. These changes in copper coordination in prion resistant and susceptible species provide new insights into the molecular mechanisms governing the resistance or susceptibility of certain species to TSE.

## INTRODUCTION

Prion diseases or transmissible spongiform encephalopathies (TSE) are neurodegenerative disorders affecting humans and animals. TSE are caused by the misfolding of the α-helical form of the physiological cellular prion protein (PrP^C^) into a β-sheet rich isoform called prion or PrP^Sc^ (1). TSE are rare disorders that can be sporadic, genetic or infectious. Animal TSE include scrapie in sheep and goats, chronic wasting disease (CWD) in cervids and bovine spongiform encephalopathy (BSE) in cattle (2).

The PrP^C^ structure consists in a C-terminal folded domain (from residues 128 to 231, hereafter in human numbering) mainly containing α-helical motifs and two short anti-parallel β-sheets (3). On the contrary, the N-terminal moiety (residues 23-127) is largely unstructured (4) and features an octapeptide-repeat region (OR) (residues 61-91) composed by four octapeptides, each carrying histidines able to coordinate prevalently one copper ion, Cu(II) (5). Cu(II) can also bind at two additional histidines -H96 and H111 with coordination from the imidazole rings and nearby backbone amides-located in a segment (residues 90-111) called “fifth” or non-OR copper binding site (6) (Figure 2A). Adjacent to the non-OR region is the palindromic motif of sequence AGAAAAGA (residues 113-120) known to be able to initiate neurotoxic β-sheet formation (7–9). Although PrP^C^ and PrP^Sc^ have identical primary sequence, they have distinct physicochemical properties. PrP^C^ exists as a detergent-soluble monomer and is readily degraded by proteinase K (PK), whereas PrP^Sc^ forms detergent-insoluble aggregates and shows high resistance to PK digestion (10). Following treatment with PK, PrP^Sc^ typically generates a protease-resistant core, referred to as PrP27-30, which is N-terminally truncated at around residue 78 (11) and it is sufficient to support prion replication and disease progression (12).

**Figure 1.**
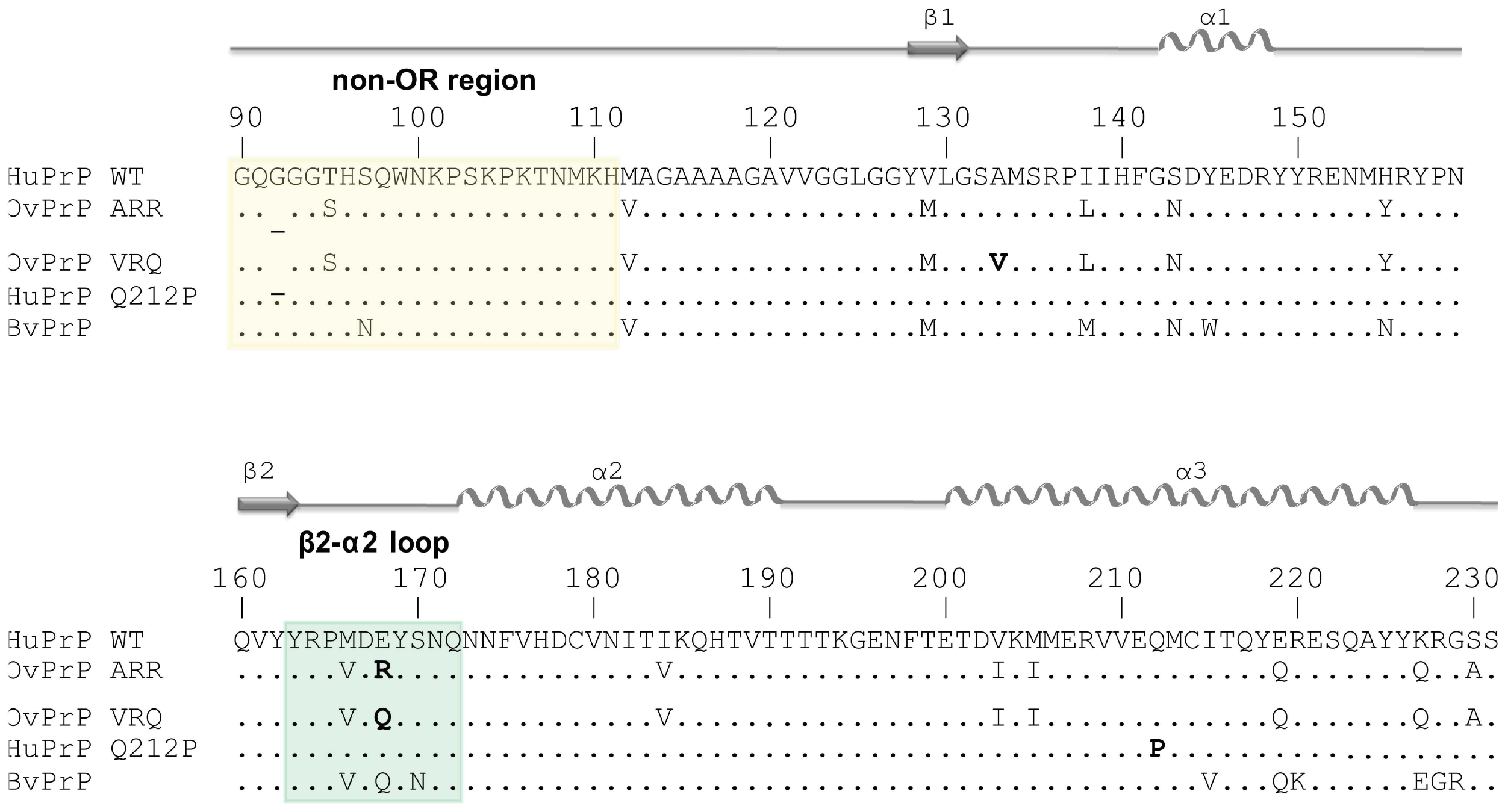
Amino acid sequences and secondary prion protein structure of HuPrP WT, OvPrP ARR, OvPrP VRQ, HuPrP Q212P and BvPrP. Comparison of amino acid sequences and secondary prion protein structure of human (HuPrP; *Homo sapiens sapiens*, GenBank accession number AAH22532), ovine with the polymorphic residue position (Q/R) (OvPrP; *Ovis aries*, AFM91142) and bank vole (BvPrP; *Myodes glareolus*, AAL57231). On the top, the secondary structure elements are shown. The yellow box highlights the non-OR copper binding site (residues 90-111) and the green box the β_2_-α_2_ loop (residues 163-172).

**Figure 2.**
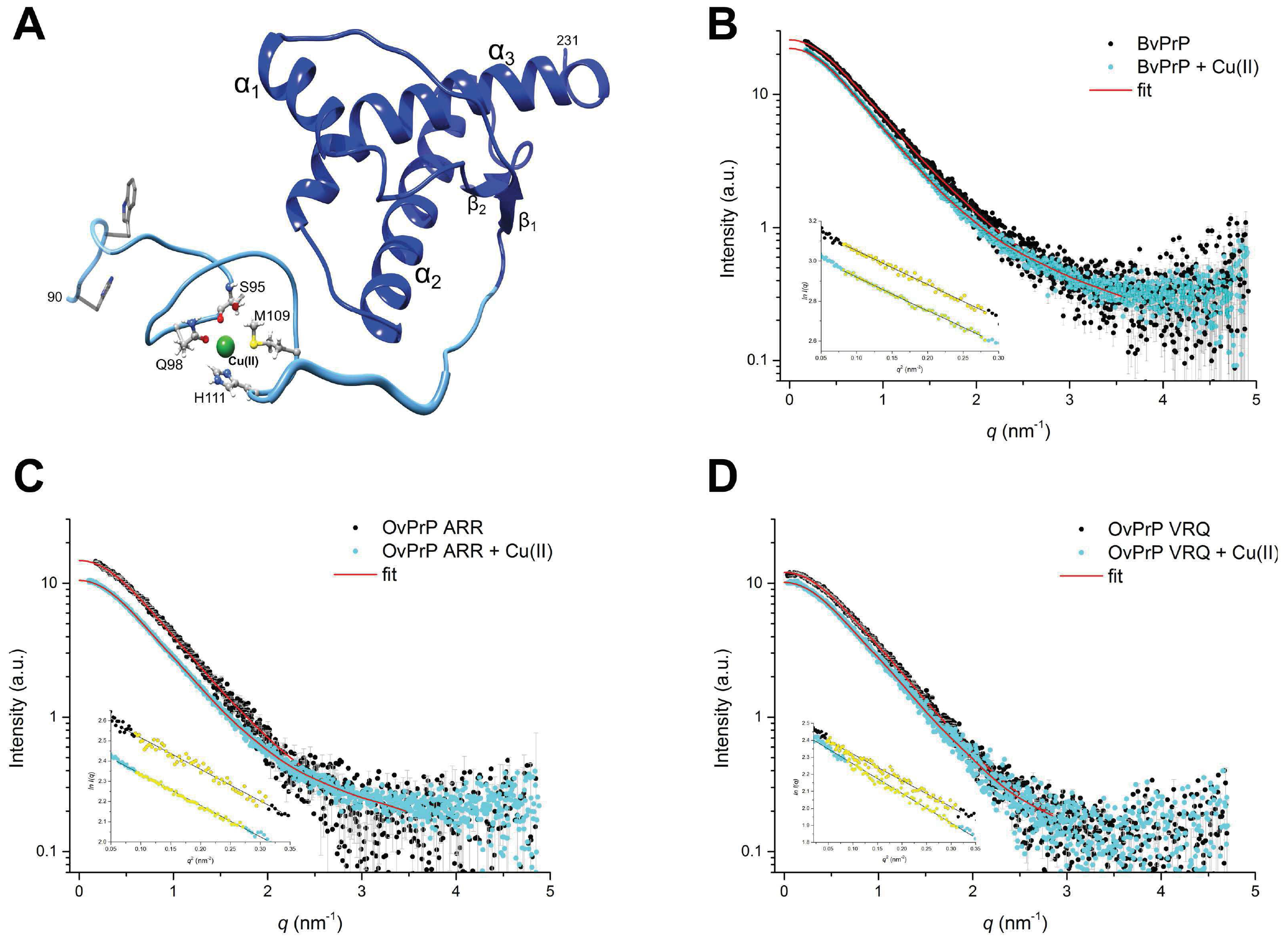
SAXS measurements on *apo* and Cu(II)-recPrP. (**A**) Cartoon representation of the C-terminal HuPrP with the non-OR copper binding site. The structure of the non-OR copper binding site from residue 90 to 126 is shown in blue; in ball and stick the residues coordinating a copper ion (S95, H96, Q98, H111, M109, in human numbering), the copper ion is shown as a dark green sphere. The palindromic region is shown with an enlarged ribbon. (**B-D**) SAXS curves of BvPrP, OvPrP ARR and OvPrP VRQ, respectively. Black and light blue dots represent the *apo* and Cu(II)-recPrP SAXS curves, respectively, with in red the GNOM fitting. Insets show the Guinier fits (yellow dots).

The functional implications of Cu(II)-binding to PrP^C^ are not unequivocal. Compelling evidences propose a Cu(II)-mediated neuroprotective role for PrP^C^ functions as modulator of synaptic plasticity and S-nitrosylation (13–14); some others point out the role of Cu(II) either as promoter or attenuator of β-sheet conversion and amyloidal aggregation (15–18). The proximity of the non-OR region to the amyloid core suggests a possible link between Cu(II) binding and prion conversion (15, 19). We have recently reported that pathogenic PrP^C^ genetic mutations affect Cu(II) coordination in the non-OR region and this altered coordination promotes prion conversion *in vitro* and in cellular models (20–21). The involvement of H96 and H111 in non-OR region shows that Cu(II) occupancy plays a role in determining the conformation of this section inducing novel long-range interactions between the N- and C-terminal PrP^C^ region with possible physiological significance in prion conversion (16). Given the importance of the C-terminal region for prion propagation and the controversial role of Cu(II) as attenuator or facilitator of TSE, further studies on the Cu(II)-PrP^C^ interactions are of pivotal importance to clarify the conformational and functional consequences of Cu(II) binding to PrP^C^.

Here, we set out to investigate the Cu(II) structural effects and its coordination geometries when bound to the non-OR region of recombinant PrP^C^ (recPrP) from animal species considered susceptible or resistant to TSE by means of small angle X-ray scattering (SAXS) and extended X-ray adsorption fine structure (EXAFS) spectroscopy. As TSE-resistant model, we used the C-terminal truncated form of ovine PrP^C^ carrying the protective polymorphism α_1_36, R154 and R171 (OvPrP ARR); while as highly TSE-susceptible PrP^C^ models we employed truncated OvPrP with polymorphism V136, R154 and Q171 (OvPrP VRQ) and Bank vole recPrP (BvPrP) (22–23) (Figure 1).

SAXS is an established method for structural characterization of biological macromolecules in solution and it is directly applicable to the study of flexible systems such as intrinsically disordered proteins and multi-domain proteins with unstructured regions like, for instance, PrP^C^ (24–25). Flexible particles are difficult objects to study and often little is known about their structural organization in native conditions. Consequently, standard high-resolution structural biology approaches -such as X-ray crystallography, NMR and cryo-electron microscopy-are limited in their ability to characterize disordered systems. Without the requirement for crystals and without effective size limitations, SAXS in near-native solutions is becoming more and more popular for the characterization of such systems, providing relevant information in terms of molecular shape and structural flexibility. In this study, SAXS provided new insights into the conformational aspects governing the interaction between Cu(II) and the non-OR region. The metal promotes a significant global compaction in all the different recPrP molecules. The reduction of SAXS dimensional parameters (*R_g_* and *D_max_*) of Cu(II)-bound recPrP indicates a decrease of local flexibility in the non-OR region possibly due to transient interactions with the C-terminal region.

Subsequently, we used EXAFS spectroscopy as sensitive technique to study the coordination geometry of Cu(II) bound to OvPrP (ARR and VRQ) and BvPrP. X-ray absorption spectroscopy is exquisitely sensitive to the coordination geometry of an absorbing atom and therefore allows the determination of bond distances and angles of the surrounding atomic cluster to be measured with near-atomic resolution (26). We identified two Cu(II) coordination geometries, namely *type-1* and *type-2*. In *type-1*, Cu(II) is bound to the side chains of four amino acids (S95, Q98, M109 and H111); this coordination is present in OvPrP ARR and in WT HuPrP. Conversely, *type-2* is present in TSE-susceptible species: bank vole, sheep with VRQ polymorphism and human with pathogenic mutations (20–21), where Cu(II) is coordinated by three amino acids (Q98, M109 and H111) and by one water molecule.

Our results reveal that Cu(II) affects the structural plasticity of the non-OR region leading to a more compacted conformation of recPrP. We also observe that the non-OR Cu(II) coordinations changes in the recPrP of TSE-resistant and susceptible species. These data support the hypothesis that amino acid variations observed in mammalian PrP^C^ sequences may have structural effects on both the globular domain and the N-terminal moiety, particularly in the non-OR region with consequences on Cu(II) coordination. These changes in copper coordination in prion resistant and susceptible species have important physiological implications, providing new insights into the molecular mechanisms governing the resistance or susceptibility of certain species to TSE.

## MATERIALS AND METHODS

### Plasmids construction, protein expression and purification

The pET-11a plasmid (Novagen) encoding for the truncated BvPrP (residues 90-231), OvPrP VRQ (residues 94-234, carrying the TSE-susceptible polymorphism V136, R154 and Q171) and OvPrP ARR (residues 94-234, carrying the TSE-resistant polymorphism A136, R154 and R171) were purchased from Genewiz (Germany, GmbH). All the recombinant proteins were expressed, purified and *in vitro* refolded according to our previous protocols (20–21).

### SEC-SAXS measurements, data analysis and modelling

All the experiments were performed at the ESRF BioSAXS beamline BM29, Grenoble, France (27). Given the sensitivity for batch SAXS mode for even small amounts of large soluble aggregates we used SEC-SAXS approaches to measure SAXS data only on monodisperse samples. A volume of 250 μL of protein *per* each BvPrP sample (*apo* and copper-loaded) at 12 mg/mL was loaded on a GE Superdex 75 10/300 GL column, and a volume of 50 μL protein *per* each OvPrP ARR and OvPrP VRQ (*apo* and copper-loaded) samples at 7 mg/mL was loaded on a GE Superdex 200 5/150 GL column *via* a high performance liquid chromatography (HPLC) system (DGU-20A5R, Shimadzu, France) attached directly to the sample-inlet valve of the BM29 sample changer (28). All the *apo* samples were measured in buffer 25 mM Sodium Acetate, 250 mM NaCl, pH 5.5 at 20 °C. Cu(II) loading on recPrP samples were achieved by dialysis (Spectra/Por 3.5 kDa MWCO membrane) against buffer 25 mM Sodium Acetate, 250 mM NaCl, pH 5.5 containing CuSO4 at 1:1 (Cu(II):recPrP) molar ratio at 4 °C for 12 hours, and then against buffer 25 mM Sodium Acetate, 250 mM NaCl, 1 μM of CuSO4, pH 5.5 to remove the excess of metal at 4 °C for 4 hours. After centrifugation (30 minutes, 16,000 g at 4 °C), Cu(II)-samples were then measured in buffer 25 mM Sodium Acetate, 250 mM NaCl, 1 μM of CuSO4, pH 5.5 at 20 °C. The columns were equilibrated with 3 column volumes to obtain a stable background signal that was confirmed before measurement. All the SAXS data were collected at a wavelength of 0.99 Å using a sample-to-detector (PILATUS 1 M, DECTRIS) distance of 2.81 m. The scattering of pure water was used to calibrate the intensity to absolute units (29). Data reduction was performed automatically using the EDNA pipeline (30). Frames in regions of stable *R_g_* were selected and averaged using PRIMUS (31) to yield a single averaged frame corresponding to the scattering of individual SEC species. All parameters for SAXS analysis, sample details and results are described in Table S1 according to recent recommended guidelines (32). Briefly, analysis of the overall parameters was carried out by PRIMUS from ATSAS 2.8.4 package (33) and by ScÅtter 3.0 software. The pair distance distribution function, P(r), and maximum diameter of the particle (*D_max_*) were calculated in GNOM using indirect Fourier transform method (34). Protein molecular masses were estimated using both Porod volume (34) and scattering mass contrast (32) methods. For low-resolution structural models, Ensemble Optimization Method (EOM) modeling was conducted using models of the BvPrP (amino acids 170-231, PDB id 2K56) and OvPrP (amino acids 170-231, PDB id 1Y2S) previously solved by NMR (35–36), and the rest of the protein was represented as beads corresponding to individual residues (37). EOM employs a genetic algorithm to select subsets of conformations from the random pool that best fits the experimental data. The selected ensembles represent a low-resolution sample space used to generate distributions of structural parameters (37–38). An initial random pool of 10,000 models was generated in RanCh (version 2.1) (38). Final ensembles were selected from the starting pool using a genetic algorithm implemented in GAJOE (version 2.1) (38). All SAXS data were deposited into SASBDB data bank (ID: SASDEW7, SASDEX7, SASDEY7, SASDEZ7, SASDE28, SASDE38, see Table S1).

### XAS spectra measurements and data analysis

Samples with 1:1 (Cu(II):recPrP) molar ratio were prepared in 25 mM NaOAc, pH 5.5, with a protein concentration of ~1 mM. Briefly, recPrP were first dialyzed (Spectra/Por 3.5 kDa MWCO membrane) against buffer containing 25 mM NaOAc, 1 mM of CuSO4, pH 5.5 and then against the same buffer containing 1 μM of CuSO_4_ to remove the excess of unbound metal. Sample monodispersity after Cu(II) loading was assessed by SEC using a GE Superdex 200 Increase 10/300 GL column. X-ray absorption spectra were recorded at ESRF on BM30B FAME beamline (39). The spectra were collected at the Cu K-edge in fluorescence mode using a solid state 30-element Ge detector, with sample orientation at 45° to incident beam. The X-ray photon beam was vertically focused by a Ni-Pt mirror, and dynamically, sagittally focused in the horizontal size. The monochromator was equipped with a Si(111) double crystal, in which the second crystal was elastically bent to a cylindrical cross section. The energy resolution at the Cu K-edge is 0.5 eV. The spectra were calibrated by assigning the first inflection point of the Cu foil spectrum to 8981 eV. All the spectra were collected at 10 K. For Cu(II) samples, photo reduction is usually observed and thus the beam was moved to different spots of the sample at each scan. During collection, data were continuously monitored in order to insure sample homogeneity across the multiple spots collected from different sample-holder’s cells. The following samples were measured: Cu(II)-OvPrP ARR, Cu(II)-OvPrP VRQ and Cu(II) BvPrP. For each sample, 12 spectra were recorded with a 7 s/point collection statistic and averaged. The collection time was 25 min for each spectrum.

The analysis of the EXAFS data was carried out using the GNXAS code (40–41) which is based on a theoretical calculation of the X-ray absorption fine structure signal and a subsequent refinement of the structural parameters. In the GNXAS approach the interpretation of the experimental data is based on the decomposition of the EXAFS *χ*(k) signal into a summation over n-body distribution functions *γ*^(n)^, calculated by means of the multiple scattering (MS) theory. Each signal has been calculated in the muffin-tin approximation using the Hedin-Lundqvist energy dependent exchange and correlation potential model, which includes inelastic loss effects. The analysis of the EXAFS spectra was carried out starting from the coordination models reported in the literature for the WT HuPrP and HuPrP Q212P proteins (6, 20-21), and considering the amino acid sequences of the species. In particular, the analysis of the OvPrP ARR resistant specie has been carried out considering the coordination with a nitrogen atom of H111, with two oxygen atoms of S95 that chelates the Cu(II) ion forming a ring in the equatorial pane (in this case two carbon atoms of the serine give rise to a single scattering contribution at about 2.86 Å), with an oxygen atom of Q98 and a sulfur atom of M109. The EXAFS spectra of the more susceptible OvPrP VRQ and BvPrP species have been analysed using the same model as HuPrP Q212P where the Cu(II) ion is coordinated by H111, Q98, M109 and a water molecule. Based on these two models theoretical EXAFS spectra were calculated to include contributions from first shell two-body signals and many body configurations. Previous investigations on model compounds have shown that a quantitative EXAFS analysis of systems containing histidine rings or having amino acid residues that are chelated to the Cu(II) ion, requires a proper treatment of MS contributions (6, 20-21, 42). In particular, the EXAFS analysis of systems containing histidine rings requires a proper treatment of MS four-body terms associated with the Cu-N-C-C(N) configurations, while coordination with S95 gives rise to three-body terms associated with the Cu-O-C configuration having a multiplicity of two. The structural parameters used in the fits are the bond distance (*R*) and bond variance (*σ^2^_R_*) for a two-body signal, the two shorter bond distances, the intervening angle (*θ*) and the six covariance matrix elements for a three-body signal. The four-body configurations are described by six geometrical parameters, namely, the three bond distances, two intervening angles (*θ* and *φ*), and the dihedral angle (*Ψ*) defining the spatial orientation of the three bonds. These parameters were allowed to float within a preset range, typically ± 0.05 Å and ± 5° for distances and angles respectively. During the minimization procedures, the magnitudes of the Debye-Waller terms were assumed to increase with distance, and atoms at similar distances from the copper ion were assigned the same value. In all cases two additional nonstructural parameters are minimized, namely E_0_ (core ionization threshold) and S_0_^2^ (many body amplitude reduction factor). To establish error limits on the structural parameters, a number of selected parameters from the fit results are statistically analyzed using two-dimensional contour plots. This analysis examines correlations among fitting parameters and evaluates statistical errors in the determination of the copper coordination structure, as previously described (20). Briefly, parameters with highest correlation dominate in the error estimate. The quality of the fits is determined by the goodness-of-fit parameter, *R_i_* (42), and by careful inspection of the EXAFS residuals.

## RESULTS

### Mammalian prion proteins undergo major compactness changes in presence of copper

The structural consequences of Cu(II) on PrP^C^ were previously investigated by SAXS on full-length murine recPrP (MoPrP) reporting a global compaction of the protein due to inter-domain interactions upon metal binding to the eight tandem repeat region (16). Here, we investigated by SAXS the Cu(II) role on the solution structures of C-terminal truncated recPrP carrying only the non-OR region as metal binding site. Structural differences due to Cu(II) interaction with the flexible N-terminal moiety (residues 90-127) may provide new insights into the molecular determinants governing the different TSE susceptibility observed in OvPrP ARR (TSE resistant), in OvPrP VRQ and BvPrP (TSE susceptible). The *R_g_*, I(0) and UV traces as functions of frames show that recPrP were highly pure and well separated in individual peaks (Figure S1). RecPrP samples remained monodispersed after Cu(II) loading (Figure S2). All SAXS results are exposed in Table S1. Data frames under each of the main elution peaks -for which the *R_g_* values were the same within error and statistically indistinguishable as assessed using CorMap (43)-were selected and averaged for further analysis. Primary data analysis from scattering curves showed that both *apo* and Cu(II)-recPrP are very similar with an elongated and flexible shape (Figure 2, B-D) as previously observed for the *apo* and Cu(II)-loaded full-length MoPrP (16). Similar conclusions regarding flexibility can be drawn from Kratky plots (Figure S3). The *R_g_* of recPrP determined by Guinier analysis showed small differences between *apo* and Cu(II)-bound proteins (Figure 2, B-D *insets*). In particular, the calculated *R_g_* of *apo* BvPrP (2.41 nm), OvPrP ARR (2.32 nm) and OvPrP VRQ (2.31 nm) were slightly larger than the *R_g_* of the same Cu(II)-bound recPrP (2.37 nm, 2.25 nm and 2.24 nm, respectively). For comparison, the *R_g_* of BvPrP(119-231) and OvPrP(119-231) calculated from the solution NMR structures (35–36) were 1.65 and 1.69 nm, respectively; however, these structures lack atomic coordinates for residues 90 to 118. Overall, the molecular dimensions observed for the *apo* recPrP are in agreement with previous SEC-SAXS studies on MoPrP(89-230) (44). Distance distribution function, P(r) analysis, revealed reduction in the *D_max_* values from ~9.4 nm for *apo* recPrP to ~8.7 nm for Cu(II)-recPrP. For all the proteins, the *R_g_* and I(0)-based mass values were in excellent agreement with the expected monomeric recPrP(90-231) molecular weight (*i.e.* ~16 kDa, see Table S1).

The N-terminal region (residues 90-127) of C-terminal recPrP is largely flexible in solution (4). Hence, we analyzed the data using EOM, which gives useful information such as *R_g_* and *D_max_* distributions in case of proteins with flexible domains. EOM analysis of recPrP yielded good quality fits for the *apo* and Cu(II)-proteins (Figure 3 *insets* and Table S1). Size distributions (*R_g_*) of *apo versus* Cu(II)-recPrP provided qualitative assessment on the structural effect of metal to protein compactness through direct comparison of the distributions of the selected ensembles and the pool (Figure 3 A-C,). The EOM size distributions showed multimodal distributions that converge into a major population with *R_g_* and *D_max_* of ~2.2 nm and ~8.3 nm, respectively, for the *apo* proteins, and with *R_g_* and *D_max_* of ~2.0 nm and ~7.8 nm, respectively, for Cu(II)-recPrP. Interestingly, the ensemble conformers of TSE-resistant OvPrP ARR loaded with Cu(II) display major reduction of structural parameters (*i.e. R_g_* 1.93 nm and *D_max_* 6.57 nm) compared to TSE-susceptible Cu(II)-OvPrP VRQ and Cu(II)-BvPrP. The results from the EOM analysis of the size distribution are in agreement with values obtained from P(r) distribution function, where a reduction of ~0.6 nm has been observed for *D_max_*of Cu(II)-protein compared with the *apo* form. Our results indicate that a significant amount of compaction of the extended conformation of *apo* recPrP occurs upon Cu(II) binding (Figure 3, D). Previous studies interpreted the reduction in the *R_g_* and *D_max_* leading to the global compactness of full-length Cu(II)-MoPrP, as result of a decrease in the flexibility of the N-terminal region, which exhibits interaction with the C-terminal globular domain upon metal binding (16). Here, we quantitatively measured the protein flexibility using two metrics, *R_flex_* and *R_σ_*, available in the EOM analysis, thus complementing the low-resolution structural descriptors (38). Using *R_flex_* metric, the selected ensemble distributions can be numerically compared to that of the pool, the latter representing a reference for flexibility. For instance, the quantification of the flexibility of *apo* OvPrP ARR (ensemble *R_flex_* = 82.62% *versus* pool *R_flex_* = 84.37%) and Cu(II)-OvPrP ARR (ensemble *R_flex_* = 80.73% *versus* pool *R_flex_* = 86.42%) confirm numerically the effect of copper on the flexibility of the protein. Both *apo* and Cu(II)-recPrP are flexible systems, with *R_flex_* values of ~80%, but show less flexibility in the presence of copper as compared to the threshold of flexibility computed from the Cu(II)-recPrP pools, *i.e.* ~88% (Table S1 *panel e*).

**Figure 3.**
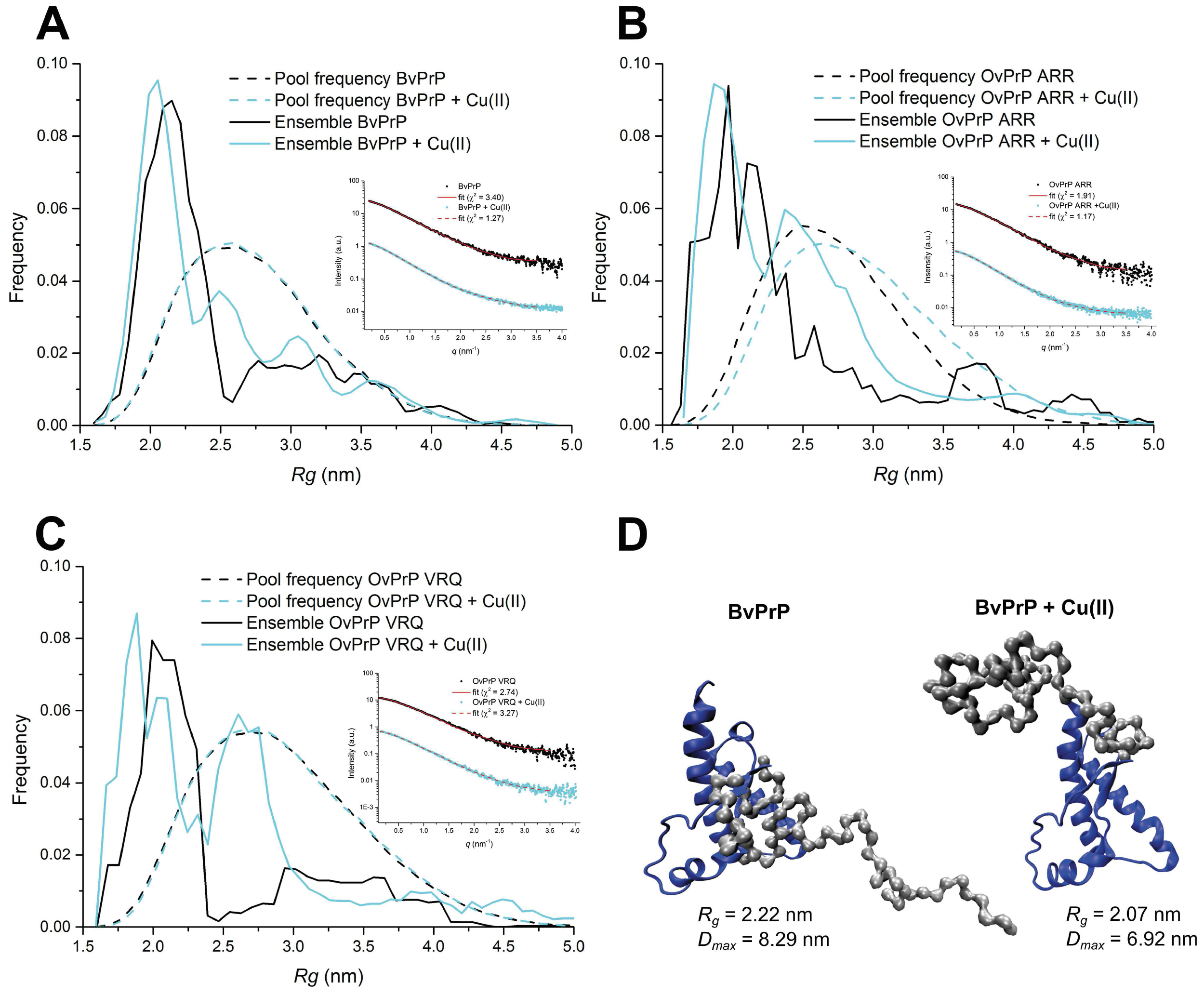
Characterization of the flexibility of *apo* and Cu(II)-bound recPrP using EOM. (**A-C**) Size distributions (*R_g_*) of BvPrP, OvPrP ARR and OvPrP VRQ, respectively, providing qualitative assessment through direct comparison of the distributions of the selected ensembles (black and light blue lines for *apo* and Cu(II)-recPrP) and the pool (dotted black and light blue lines for *apo* and Cu(II)-recPrP). In the insets, I(q) *versus q* experimental SAXS profiles (black dots for *apo* proteins and light-blue dots for copper-loaded proteins) with the EOM fit models (continuum red lines for *apo* proteins and dotted lines for copper-loaded proteins with the corresponding *χ*^2^ values) of the recPrP. The curves are shifted by an arbitrary offset for better comparison. (**D**) Representation of models of *apo* and Cu(II)-BvPrP obtained with EOM with the most representative structural parameters (*R_g_* and *D_max_*) from the ensembles.

### Copper coordination in the non-OR region of prion resistant species

The Cu(II) coordination structure in the non-OR binding site of HuPrP WT was unambiguously assessed by EXAFS in previous investigations (6, 20). The Cu(II) was found to be coordinated by two histidines (H96 and H111) and by Q98 and M109. It was proposed that the non-OR region is stabilized when the Cu(II) ion is coordinated by H96 and H111, and this coordination prevents prion conversion (21). The EXAFS experimental spectrum of HuPrP WT extracted with o 1 a three-segmented cubic spline shows a typical feature around *k* = 5 Å^-1^ (Figure 4 A) while the Fourier Transform (FT) spectrum is characterized by a first-shell peak centered at 1.5 Å and additional high intensity outer shell peaks in the distance range between 2 and 4 Å (Figure 4 C) that are indicative of two histidines coordinated to the metal ion (6, 20–21). The H96 residue is present in the OvPrP ARR amino acid sequence but the EXAFS spectrum of this species is different from that of HuPrP WT showing markedly different features in the k range around 5 Å^-1^, that is sensitive to the His ligands (Figure 4 A, B). Additionally, the intensity of the second peak of the FT at 2.2 Å of OvPrP ARR is lower as compared to HuPrP WT (Figure 4 C, D) as only one histidine ligand coordinates the copper center, but the overall shape and intensity of the FT higher distances peaks indicate that an additional amino acid is chelated to Cu(II) ion (42).

**Figure 4.**
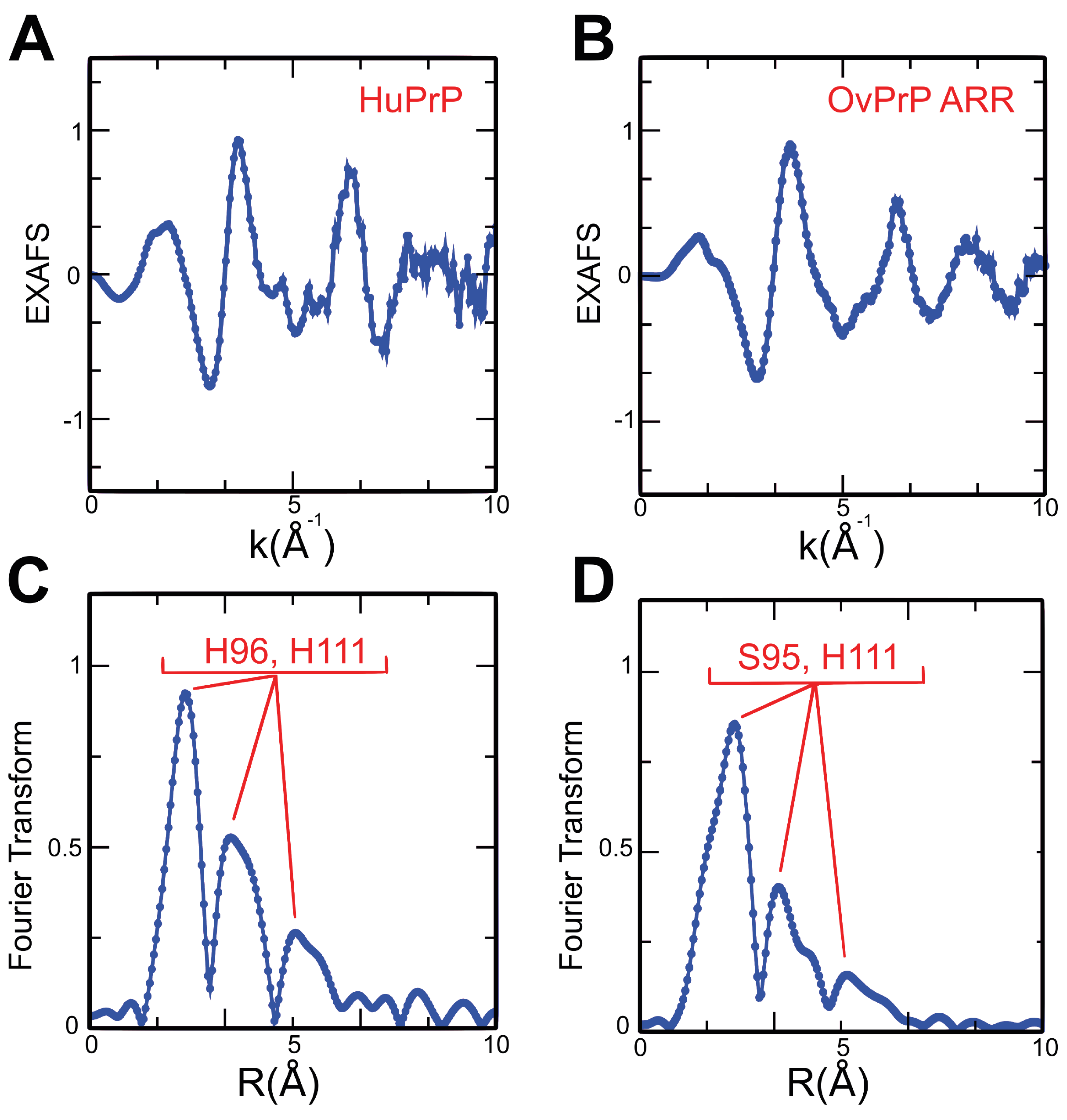
Cu K-edge EXAFS experimental spectra extracted with a three-segmented cubic spline of (**A**) Cu(II)-HuPrP WT [data from reference (20)], (**B**) Cu(II)-OvPrP ARR. Non phase shift-corrected Fourier transforms of the EXAFS experimental spectrum calculated in the interval k = o 1 2.1-10.0 Å^-1^ of (**C**) Cu(II)-HuPrP WT, and (**D**) Cu(II)-OvPrP ARR.

By comparison of the amino acid sequences of HuPrP and OvPrP ARR (Figure 1) in the region between residues 92 and 96, two serine (S95 and S97) are present in these recPrP and both can chelate Cu(II) (42). Here, we assigned S95 as the ligand of Cu(II) ion. Starting from these observations a quantitative analysis of the EXAFS data of OvPrP ARR has been carried out using a coordination model around the Cu(II) ion comprising S95, H111 and Q98 in the equatorial plane. In particular, S95 is chelated to Cu(II) ion through two oxygen atoms forming a 6-fold ring (42) and this gives rise to two Cu-O-C three-body configurations that have to be accounted for in the analysis of the EXAFS spectra. Moreover, two second shell Cu-C contributions are present at a distance of about 2.86 Å. Additional MS contributions are associated with the H111 ring and two Cu-N-C-C(N) four-body contributions have been considered. A theoretical *χ*(k) signal has been calculated including all the relevant two-body and MS contributions and a fitting procedure has been carried out in the *k* range between 2.4 and o 1 12.5 Å^-1^ in order to find the set of structural parameters that provides the best agreement with the experimental data. In the analysis the C-N and C-O of the amino acid residues are kept fixed to 1.36 and 1.34 Å, respectively. The results of the minimization procedures are shown in Figure 6 A for OvPrP ARR. From the top of Figure 6 A the following theoretical signals are shown: the two-body contribution associated with the two S95 and one Q98 Cu-O first shell distances, the two-body contribution associated with the H111 Cu-N first shell distance, the two-body contribution associated with the M109 Cu-S first shell distance, the two-body contribution associated with the two second shell Cu-C distances of S95 residue, the two-body contribution associated with the Cu-O water distance, the MS contributions associated with two Cu-O-C three-body configurations of the S95 residue, the MS contribution associated with two Cu-N-C-C(N) four-body configurations of the H111 residue, and the total *χ*(k) signal compared with the experimental spectrum. The EXAFS theoretical signals match the experimental data quite well, thus confirming the validity of the Cu(II) coordination model proposed on the basis of the EXAFS, FT characteristic fingerprints as well as on the amino acid sequences. The structural parameters obtained from the EXAFS analysis are listed in Table 1. In OvPrP ARR S_0_ was found equal to 0.9, while E0 was found 3 eV above the first inflection point of the spectra. The results of the fitting procedure confirm that copper is coordinated with H111 (Cu-N distance is 1. 98(2) Å), with S95 through two oxygen atoms (Cu-O distance 2.00(2) Å, with an additional O/N atom at 2.00(2) Å that may belong to Q98, and one sulfur scatterer at longer distance (3.23(4)/3.27(4) Å). In the final fit one oxygen atom of the solvent at 2.28(5)/2.32(5) Å was included as suggested in previous investigations (6, 20, 45). The additional oxygen improves the quality of the fit (*R_i_* improved by 20%). Note that the inclusion of the MS signal associated with S95 is essential to reproduce the experimental spectrum (*R_i_* improved by 30%) and this strongly supports the validity of the proposed coordination model.

**Table 1.** Structural parameters derived from the EXAFS analysis. Structural parameters determined from the fit of the EXAFS data at the Cu K-edge of Cu(II)-OvPrP ARR, Cu(II)-OvPrP VRQ, and Cu(II)-BvPrP. *N* is the coordination number, *R* is the distance between the copper ion and the ligand, *σ*^2^is the Debye-Waller factor. Statistical errors are reported in parentheses.

| **OvPrP ARR** |  |  | **OvPrP VRQ** |  |  | **BvPrP** |  |  |
| --- | --- | --- | --- | --- | --- | --- | --- | --- |
| *N* | *R* (Å) | $\sigma^2$ (Å<sup>2</sup>) | *N* | *R* (Å) | $\sigma^2$ (Å<sup>2</sup>) | *N* | *R* (Å) | $\sigma^2$ (Å<sup>2</sup>) |
| 1 N<sub>His</sub> | 1.98(2) | 0.008(3) | 1 N<sub>His</sub> | 2.00(2) | 0.008(3) | 1 N<sub>His</sub> | 2.00(2) | 0.008(3) |
| 3 O/N | 2.00(2) | 0.006(3) | 3 O/N | 1.99(3) | 0.009(3) | 3 O/N | 1.99(3) | 0.009(3) |
| 1 O | 2.32(5) | 0.014(4) | 1 O | 2.41(3) | 0.010(4) | 1 O | 2.43(3) | 0.009(4) |
| 1 S | 3.27(4) | 0.009(4) | 1 S | 3.26(4) | 0.015(4) | 1 S | 3.23(4) | 0.013(4) |
| 2 C | 2.87(4) | 0.006(4) |  |  |  |  |  |  |

In conclusion for OvPrP ARR the EXAFS analysis reveals the existence of a distorted octahedral geometry of the copper center and we denoted this Cu(II) coordination geometry as *type-1*.

### Copper coordination in the non-OR region of prion susceptible species

In previous works we highlighted that HuPrP point mutations associated with genetic forms of prion diseases induce a dramatic modification of the non-OR binding site (20–21). In HuPrP Q212P, the non-OR binding site becomes less structured because H111, Q98, M109 and a water molecule bind to the metal. Both EXAFS and FT spectra of HuPrP Q212P are different from those of OvPrP ARR here analyzed (Figure 5 A, D). In particular, the HuPrP Q212P FT is characterized by outer shell peaks in the range between 2 and 4 Å with lower amplitude and this shows that only one histidine coordinates the Cu(II) ion. In the present work the copper coordination of two susceptible mammalian species, namely OvPrP VRQ and BvPrP, has been investigated by means of XAS. The EXAFS and FT experimental spectra of these systems are similar to those of HuPrP Q212P (Figure 5) and the low intensity of the second peak of the FT suggests that only one histidine coordinates the Cu(II) ion. Notably, in the case of TSE-susceptible mammalian species here investigated the intensity of the FT peak at 2.2 Å is lower than that of the TSE-resistant species, thus indicating that no other amino acid residues are chelated to the Cu(II) ion and the only MS contributions are those associated with H111 as in the case of HuPrP Q212P. The EXAFS data analyses of OvPrP VRQ and BvPrP have been carried out using the coordination geometry previously determined for HuPrP Q212P (20). In particular, the EXAFS data have been analyzed considering a fourfold coordination around the Cu(II) ion with His111 and three O/N scatterers in the equatorial plane. Also in this case the copper center interacts with the sulfur atom of M109 at longer distance. Starting from this model a theoretical *χ*(k) signal has been calculated and a fitting procedure has been carried out in the *k* range between 2.4 and 12.5 Å^-1^. The results of the fitting procedures are shown in Figure 6 B-C: from the top, the following theoretical signals are shown: the two-body contribution associated with three O/N atoms, the two-body contribution associated with the H111 Cu-N first shell distance, the two-body contribution associated with the M109 Cu-S first shell distance, the two-body contribution associated with the water Cu-O distance, the MS contribution associated with two Cu-N-C-C(N) four-body configurations of the H111 residue, and the total *χ*(k) signal compared with the experimental spectrum. The EXAFS theoretical and experimental curve show a very good agreement both for OvPrP VRQ and BvPrP, thus confirming the close similarity of the Cu(II) geometry between the susceptible mammalian species and HuPrP Q212P. The structural parameters obtained from the EXAFS analysis are listed in Table 1 and they are almost identical for OvPrP VRQ and BvPrP, and also in this case S_0_^2^ was found equal to 0.9, while E0 was found 3 eV above the first inflection point of the spectra. The results of the fitting procedure show that copper is coordinated with H111 (Cu-N distance is 2.00(2) Å), with three O/N atoms at 1.99(3) Å, and one sulfur scatterer at longer distance (3.26(4)/3.23(4) Å). In the final fit one oxygen atom of the solvent at 2.41(3)/2.43(3) Å was included as suggested in previous investigations (20–21), and its inclusion improves ***R_i_*** by 20%. This coordination geometry is denoted as *type-2*.

**Figure 5.**
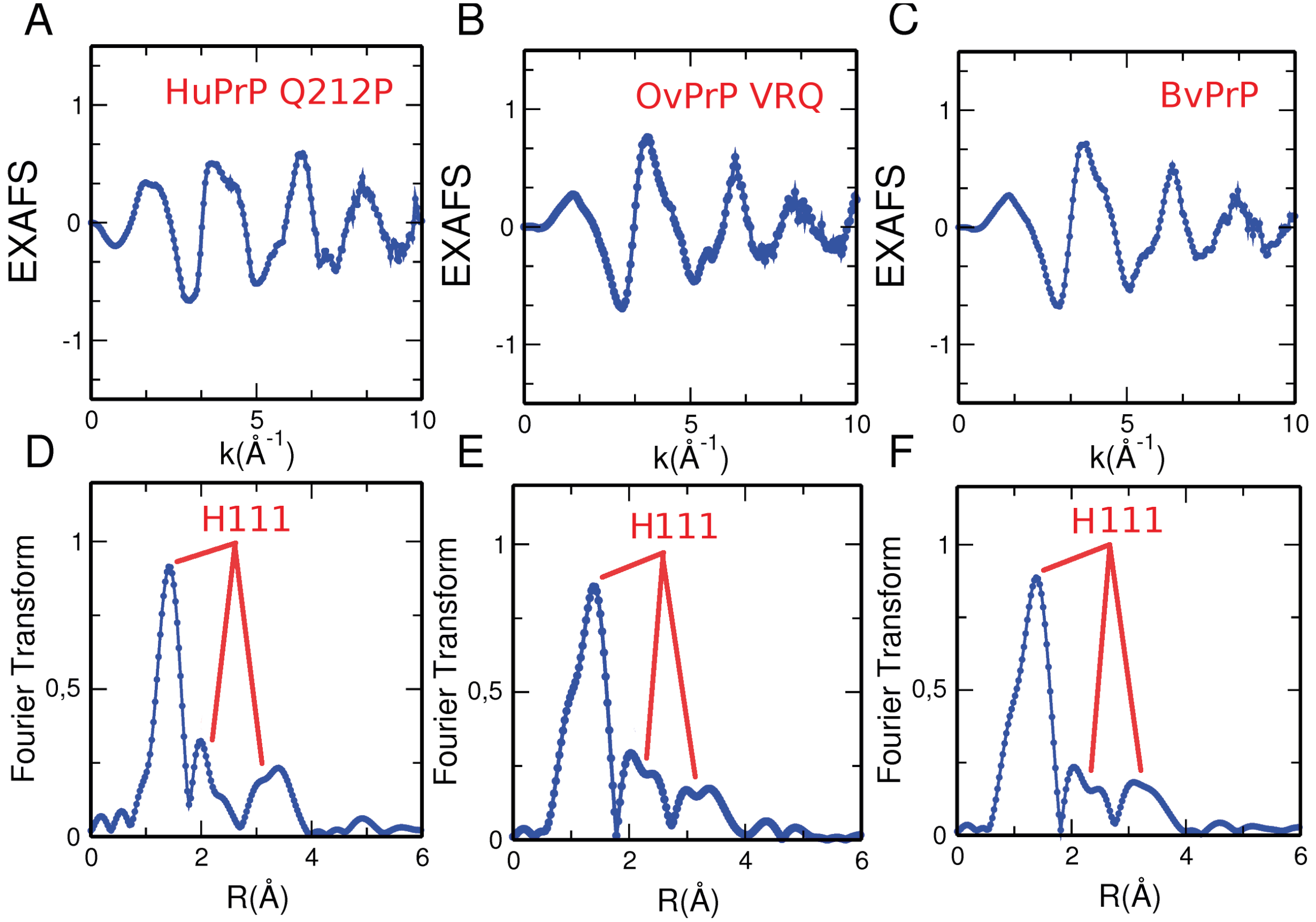
Cu K-edge EXAFS experimental spectra extracted with a three-segmented cubic spline of (**A**) Cu(II)-HuPrP Q212P [data from reference (20)], (**B**) Cu(II)-OvPrP VRQ and (**C**) Cu(II)-BvPrP. Non phase shift-corrected Fourier transforms of the EXAFS experimental spectrum o 1 calculated in the interval k = 2.1-10.0 Å^-1^ of (**D**) Cu(II)-HuPrP Q212P, (**E**) Cu(II)-OvPrP VRQ and (**F**) Cu(II)-BvPrP.

**Figure 6.**
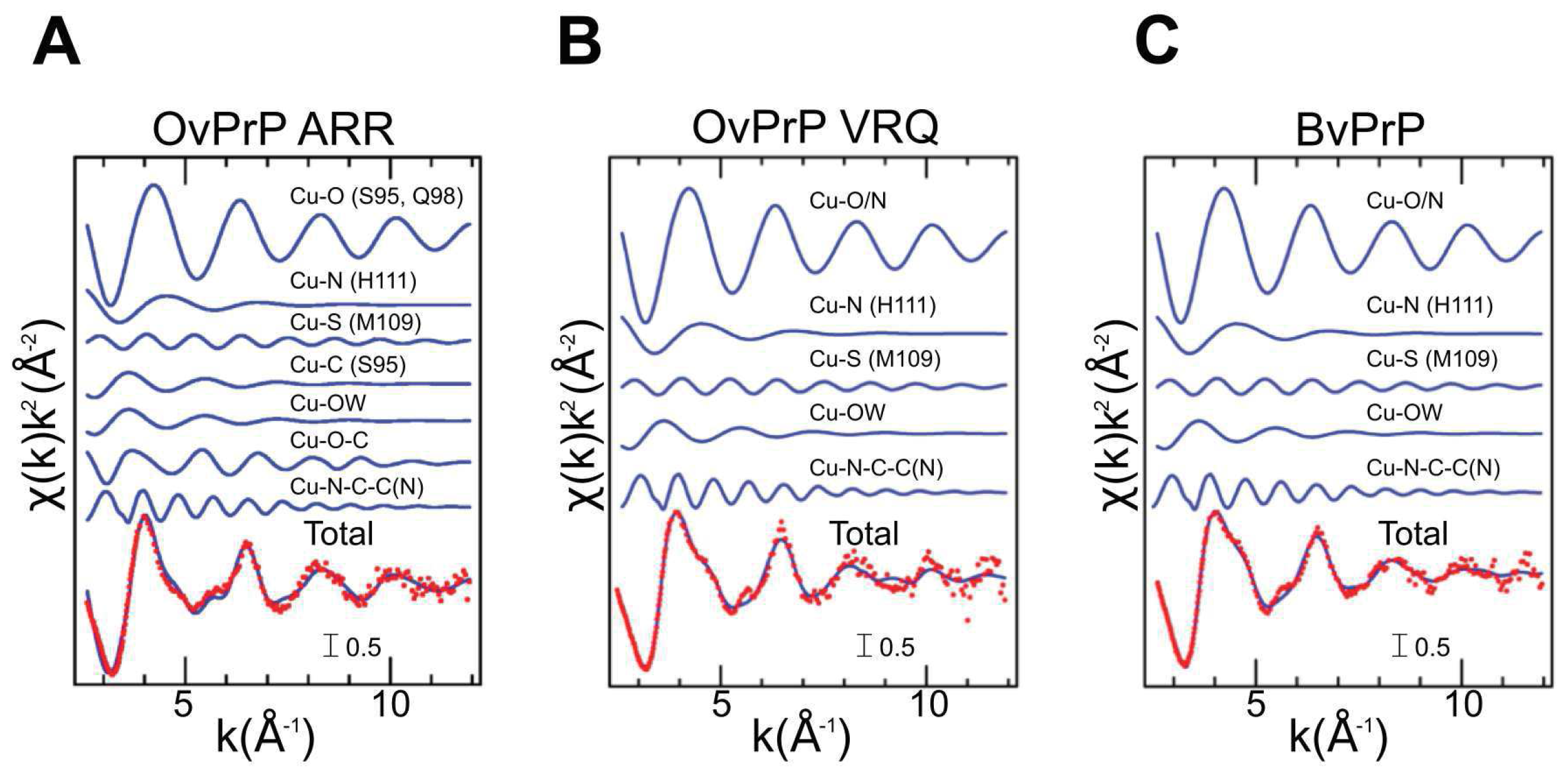
Fit of the Cu K-edge EXAFS spectrum of Cu(II)-OvPrP ARR (**A**) carried out in the −1 interval k= 2.4-12.5 Å^-1^. From the top to the bottom the following curves are reported: the Cu-O two-body theoretical signal associated with the two S95 and one Q98 oxygen atoms, the Cu-N two-body theoretical signal associated with the H111 nitrogen atom, the Cu-S two-body theoretical signal associated with the M109 sulfur atom, the Cu-C two-body theoretical signal associated with the S95 second shell carbon atoms, the Cu-O two-body theoretical signal associated with the water molecule, the Cu-O-C three-body theoretical signal associated with the S95 residue, the Cu-N-C-C(N) four-body theoretical signal associated with the H111 residue, and the total theoretical χ(k) signal (red dotted line) compared with the experimental spectrum (blue line). In panels **B** and **C** the fitting of the Cu K-edge EXAFS spectrum of Cu(II)-OvPrP VRQ and Cu(II)-BvPrP, respectively.

## DISCUSSION

PrP^C^ interacts with copper through the OR and non-OR binding sites. Different findings suggest a protective role for copper when bound to the OR region since the metal inhibits the *in vitro* amplification of PrP^Sc^-induced recPrP aggregation and fibrillization (46–47). The OR region exhibits high reduction potential for the Cu(II)/Cu(I) couple and can initiate reactive oxygen species-mediated β-cleavage of PrP^C^ at residue G90 (48–49). This may generate the N-terminally truncated form of PrP^C^ that takes part in the amyloid core during prion conversion (50). Given the known importance of the region from residues 90 to 231 for prion formation and the proximity of the non-OR region to the palindromic amyloidogenic motif, different studies have addressed the question whether Cu(II)-bound non-OR region has a role in prion generation and disease onset. A recent report showed that transgenic mice, TgPrP(H95G), with an amino acid replacement at residue H96 had shorter disease progression than WT control mice and classical clinical signs of TSE (51). We observed that alteration of Cu(II) coordination due to H96Y mutation causes spontaneous PrP^Sc^-like formation in neuronal cultured cells and accumulation in the acid compartments (21). At that time, we proposed a model whereby HuPrP coordinating copper with two histidines (H96 and H111) in the non-OR region is more resistant to prion conversion compared to the protein coordinating Cu(II) with one histidine. However, little in the way of structural information exists regarding the Cu(II)-mediated non-OR stabilization and inter-domain contacts due to the intrinsic flexible nature of the non-OR segment. Here, we provide new insights into the Cu(II) structural consequences when the metal is bound to the non-OR region, particularly with regard to recPrP from well-known animal species considered to be highly TSE-resistant (sheep expressing PrP^C^ with ARR polymorphism) or TSE-susceptible (sheep with *Prnp* gene carrying the VRQ polymorphism and the bank vole animal model) (22–23).

Using combined SAXS and EXFAS approaches we found that Cu(II) promotes significant structural compactness of recPrP upon metal binding and displays different coordination geometries when bound to TSE-resistant or TSE-susceptible recPrP. All SAXS and EXAFS measurements were carried out at pH 5.5 where the non-OR region can still coordinate one Cu(II) ion (17, 20). The capability for Cu(II) binding at acidic pH indicates that the metal could be maintained during the cycling of the protein in the acidic endosomal compartments, where PrP^C^ accumulations and prion conversion mainly occur (21, 52). Additionally, our previous experience with EXAFS spectroscopy leads us to the conclusion that remarkable differences in the Cu(II) coordination geometries are present only at acidic conditions and not at physiological pH (20–21). These data have been compared with previous measurements on Cu(II) coordinations in the non-OR region of HuPrP WT and of HuPrP mutants (Q212P and P102L) (21).

We applied the SEC-SAXS method that allowed us to collect and interpret SAXS data on experimentally difficult aggregation-prone protein system, such as truncated recPrP, and to minimize radiation damage thanks to the continuous flow (28). To dissect the different conformational recPrP states we used EOM approaches (37–38) that provided a semi-quantitative assessment on the structural effect of Cu(II) to protein compactness through direct comparison of the size distributions of the *apo versus* Cu(II)-conformers. To the best of our knowledge, this is the first SEC-SAXS investigation on recPrP with Cu(II) bound only to the non-OR region showing novel conformational aspects of Cu(II)-recPrP, such as the decrease in dimensional parameters (*R_g_* and *D_max_*) indicative of protein compaction. Then, we expanded our understanding on Cu(II) coordination in the non-OR region of different mammalian recPrP. The EXAFS results showed that in Cu(II)-OvPrP ARR the non-OR region is structured with the metal interacting with four amino acid residues (S95, Q98, M109 and H111). Conversely, Cu(II)-OvPrP VRQ and Cu(II)-BvPrP are characterized by a more flexible non-OR binding site where solvent molecules can enter and the metal has interactions with Q98, M109 and H111. The same Cu(II) coordination was previously found in HuPrP Q212P and HuPrP P102L pathological mutants (21).

Based on our results and considering the amino acid sequences of the non-OR region we proposed two models of Cu(II) coordinations (Figure 7). The *type-1* coordination displays a closed non-OR region conformation, which can be associated with TSE-resistant species likely because of higher stability of the non-OR region. In support, our EOM results on Cu(II)-OvPrP ARR indicate that this recPrP has a tendency to adopt a more compact folding than the TSE-susceptible VRQ variant and BvPrP. Instead, in *type-2* coordination a water molecule enters the coordination shell, thus leading to a less structured and solvent exposed non-OR region. We believe that this more opened conformation of the non-OR region in the *type-2* renders the overall PrP^C^ structure more flexible and prone to structural rearrangements leading to prion conversion.

**Figure 7.**
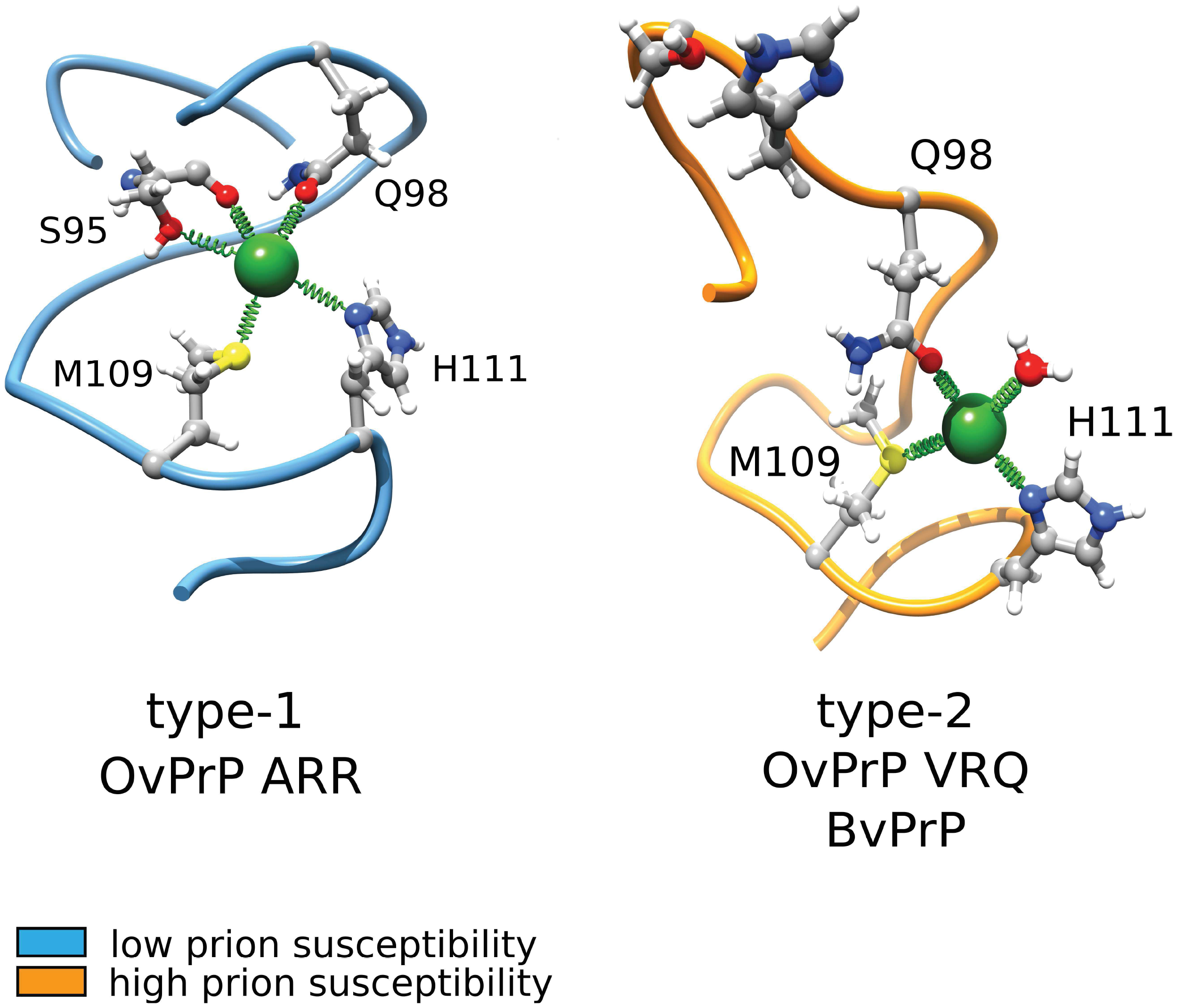
*Type-1* and *type-2* coordination models of Cu(II) in the non-OR region of TSE-resistant and TSE-susceptible species. Blue spheres identify nitrogen atoms, red spheres are oxygen atoms and yellow spheres encode for sulfur atoms. Gray and white spheres represent carbon and hydrogen atoms, respectively.

Several amino acid substitutions are present in the mammalian PrP^C^ here investigated (Figure 1). They typically affect the rigidity of some loops, through stabilizing H-bonds, and the electrostatic surface potential without affecting global folding of the structured domain. In BvPrP, the presence of N170 determines the rigidity of the β_2_-α_2_ loop (53), which is correlated with a higher susceptibility to horizontal TSE transmission and lower prion transmission barrier to interspecies transmission (54–56). In sheep, the different prion susceptibility is dictated by amino acid variations in the C-terminal globular domain. The X-ray crystal and NMR structures of the OvPrP ARR and VRQ variants reveal minor differences in the short-range H-bonding but major changes in the surface charge distribution, with ARR variant displaying surface charge variations in the α_1_-β_2_-α_2_ region (residues 154-171) (57–58).

Despite numerous endeavors in the prion field, the molecular mechanisms of the conversion remain still elusive. Structural studies on pathological point mutations have provided new clues on the early structural rearrangements occurring in some epitopes in the structured domain (45), but also revealed that amino acid variations have structural effects on the N-terminal part, affecting both Cu(II) coordination in the non-OR region (20) and long-range interactions between the Cu(II)-OR segment and the structured domain (59). Tertiary interactions between this flexible segment and the globular C-terminal domain occur when PrP^C^ binds to Cu(II), or Zn(II), through the OR region (16, 59). A recent study identified by both NMR and double electron-electron resonance EPR approaches an interacting surface made by the Cu(II)-octarepeats and a negatively charged pocket in the C-terminal domain. The Cu(II)-binding only to the OR region (authors used a full-length MoPrP H95Y/H110Y construct) perturbs globular domain residues located nearby the β_1_-α_1_ loop and the α_2_-α_3_ loop region (60). Similarly, the addition of Cu(II) to truncated MoPrP, carrying only the non-OR region, caused significant broadening of peaks corresponding in the ^15^ N-HSQC NMR spectrum to residues close to the β_1_-α_1_ loop and helix α_1_ regions (16). These observations suggest that both the Cu(II)-loaded OR and Cu(II)-non-OR regions likely bind to the same non-contiguous epitope.

Our SAXS and EXAFS observation led to the conclusion that Cu(II) in *type-1* coordination stabilizes the non-OR region and increases the protein compactness; this may favor more stable long-range interaction contacts between the 90-127 segment and the C-terminal aforementioned negatively charged pocket. Conversely, Cu(II) in *type-2* coordination renders the non-OR region more flexible and the N-terminal moiety more extended possibly due to less frequent interdomain contacts. In the *apo* form, the N-terminal moiety adopts a largely disordered conformation without stabilizing interactions with the C-terminal domain. Alterations of this inter-domain contacts may have relevant physiological implications for TSE progression since this Cu(II)-promoted non-OR *cis* interaction renders PrP^C^ more stable to PrP^Sc^ conversion. This is consistent with observations showing that an antibody, POM1, able to abolish the N- to C-terminus interaction upon binding to the negatively charged pocket causes acute neurotoxicity in mice and cultured cerebellar brain slices (61).

## CONCLUSIONS

In this work we showed that Cu(II) bound to the non-OR mediates physiologically relevant structural changes in the N-terminal moiety possibly with stabilizing interactions with the globular domain. We described two novel Cu(II) coordination geometries in the non-OR binding site of animal species considered resistant or susceptible to TSE. Changes in the non-OR conformation may dictate the lower or higher susceptibility to TSE observed in different animals. Considering the role of the non-OR and palindromic regions in supporting prion generation and propagation, this study proposes a novel structural mechanism responsible for prion susceptibility in different mammalian species.

## Author contributions

G.G., P.D. and G.L. conceived the project and jointly supervised this work. G.S., G.G., P.D. and G.L. wrote the manuscript. G.S., G.G., T.H.T., G.M. and P.D. carried out EXAFS data collection at ESRF. G.S., G.G. and P.D. analyzed EXAFS data. M.B. and G.G. carried out and analyzed SAXS data. G.S. and T.H.T. provided the recombinant protein samples. All authors read and approved the final manuscript.

## Acknowledgments

We thank andrea raspadori and paola zago for their contributions to recombinant proteins production. We are thankful to esrf for the beamtimes at bm29 and at bm 30b beamlines and to isabelle kieffer (bm 30b fame) for assistance with the measurements.

## Funding sources

This work was supported by the University of Rome “La Sapienza” (Progetto ateneo 2015, n. C26H159F5, to PD).

## Conflict of interest

The authors declare that they have no conflicts of interest with contests of this article.

## References

1. Colby DW, Prusiner SB. Prions. Cold Spring Harb Perspect Biol. 2011 Jan;3(1):a006833.

2. Imran M, Mahmood S. An overview of human prion diseases. Virol J. 2011; 8:559.

3. Surewicz WK, Apostol MI. Prion protein and its conformational conversion: a structural perspective. Top Curr Chem. 2011;305:135–67.

4. Zahn R, Liu A, Luhrs T, Riek R, von Schroetter C, Lopez Garcia F, et al. NMR solution structure of the human prion protein. Proceedings of the National Academy of Sciences of the United States of America. 2000 Jan 4;97(1):145–50.

5. Walter ED, Chattopadhyay M, Millhauser GL. The affinity of copper binding to the prion protein octarepeat domain: evidence for negative cooperativity. Biochemistry. 2006 Oct 31;45(43):13083–92.

6. Hasnain SS, Murphy LM, Strange RW, Grossmann JG, Clarke AR, Jackson GS, et al. XAFS study of the high-affinity copper-binding site of human PrP(91-231) and its low-resolution structure in solution. Journal of molecular biology. 2001 Aug 17;311(3):467–73.

7. Abskharon RN, Giachin G, Wohlkonig A, Soror SH, Pardon E, Legname G, et al. Probing the N-terminal beta-sheet conversion in the crystal structure of the human prion protein bound to a nanobody. J Am Chem Soc. 2014 Jan 22;136(3):937–44.

8. Jobling MF, Stewart LR, White AR, McLean C, Friedhuber A, Maher F, et al. The hydrophobic core sequence modulates the neurotoxic and secondary structure properties of the prion peptide 106-126. J Neurochem. 1999 Oct;73(4):1557–65.

9. Kuwata K, Matumoto T, Cheng H, Nagayama K, James TL, Roder H. NMR-detected hydrogen exchange and molecular dynamics simulations provide structural insight into fibril formation of prion protein fragment 106-126. Proceedings of the National Academy of Sciences of the United States of America. 2003 Dec 9;100(25):14790–5.

10. Oesch B, Westaway D, Wälchli M, McKinley MP, Kent SBH, Aebersold R, et al. A cellular gene encodes scrapie PrP 27-30 protein. Cell. 1985 1985/04/01/;40(4):735–46.

11. Parchi P, Zou W, Wang W, Brown P, Capellari S, Ghetti B, et al. Genetic influence on the structural variations of the abnormal prion protein. Proceedings of the National Academy of Sciences. 2000;97(18):10168–72.

12. Legname G, Baskakov IV, Nguyen HO, Riesner D, Cohen FE, DeArmond SJ, et al. Synthetic mammalian prions. Science. 2004 Jul 30;305(5684):673–6.

13. Gasperini L, Meneghetti E, Pastore B, Benetti F, Legname G. Prion protein and copper cooperatively protect neurons by modulating NMDA receptor through S-nitrosylation. Antioxid Redox Signal. 2015 Mar 20;22(9):772–84.

14. Khosravani H, Zhang Y, Tsutsui S, Hameed S, Altier C, Hamid J, et al. Prion protein attenuates excitotoxicity by inhibiting NMDA receptors. The Journal of cell biology. 2008 May 5;181(3):551–65.

15. Jones CE, Abdelraheim SR, Brown DR, Viles JH. Preferential Cu2+ coordination by His96 and His111 induces beta-sheet formation in the unstructured amyloidogenic region of the prion protein. The Journal of biological chemistry. 2004 Jul 30;279(31):32018–27.

16. Thakur AK, Srivastava AK, Srinivas V, Chary KV, Rao CM. Copper alters aggregation behavior of prion protein and induces novel interactions between its N- and C-terminal regions. The Journal of biological chemistry. 2011 Nov 4;286(44):38533–45.

17. Wells MA, Jackson GS, Jones S, Hosszu LL, Craven CJ, Clarke AR, et al. A reassessment of copper(II) binding in the full-length prion protein. Biochem J. 2006 Nov 1;399(3):435–44.

18. Wong E, Thackray AM, Bujdoso R. Copper induces increased beta-sheet content in the scrapie-susceptible ovine prion protein PrPVRQ compared with the resistant allelic variant PrPARR. Biochem J. 2004 May 15;380(Pt 1):273–82.

19. Migliorini C, Sinicropi A, Kozlowski H, Luczkowski M, Valensin D. Copper-induced structural propensities of the amyloidogenic region of human prion protein. J Biol Inorg Chem. 2014 Jun;19(4-5):635–45.

20. D’Angelo P, Della Longa S, Arcovito A, Mancini G, Zitolo A, Chillemi G, et al. Effects of the pathological Q212P mutation on human prion protein non-octarepeat copper-binding site. Biochemistry. 2012 Aug 7;51(31):6068–79.

21. Giachin G, Mai PT, Tran TH, Salzano G, Benetti F, Migliorati V, et al. The non-octarepeat copper binding site of the prion protein is a key regulator of prion conversion. Sci Rep. 2015;5:15253.

22. Goldmann W, Hunter N, Smith G, Foster J, Hope J. PrP genotype and agent effects in scrapie: change in allelic interaction with different isolates of agent in sheep, a natural host of scrapie. J Gen Virol. 1994 May;75 (Pt 5):989–95.

23. Nonno R, Bari MAD, Cardone F, Vaccari G, Fazzi P, Dell’Omo G, et al. Efficient Transmission and Characterization of Creutzfeldt-Jakob Disease Strains in Bank Voles. PLOS Pathogens. [doi:10.1371/journal.ppat.0020012]. 2006;2(2):e12.

24. Mertens HDT, Svergun DI. Structural characterization of proteins and complexes using small-angle X-ray solution scattering. J Struct Biol. 2010 Oct;172(1):128–41.

25. Bernado P, Svergun DI. Structural analysis of intrinsically disordered proteins by small-angle X-ray scattering. Mol Biosyst. 2012;8(1):151–67.

26. Holm RH, Kennepohl P, Solomon EI. Structural and Functional Aspects of Metal Sites in Biology. Chemical Reviews. [doi: 10.1021/cr9500390]. 1996 1996/01/01;96(7):2239–314.

27. Pernot P, Round A, Barrett R, De Maria Antolinos A, Gobbo A, Gordon E, et al. Upgraded ESRF BM29 beamline for SAXS on macromolecules in solution. J Synchrotron Radiat. 2013 Jul;20(Pt 4):660–4.

28. Brennich ME, Round AR, Hutin S. Online Size-exclusion and Ion-exchange Chromatography on a SAXS Beamline. J Vis Exp. 2017 Jan 5(119).

29. Orthaber D, Bergmann A, Glatter O. SAXS experiments on absolute scale with Kratky systems using water as a secondary standard. Journal of Applied Crystallography. 2000;33(2):218–25.

30. Brennich ME, Kieffer J, Bonamis G, De Maria Antolinos A, Hutin S, Pernot P, et al. Online data analysis at the ESRF bioSAXS beamline, BM29. Journal of Applied Crystallography. 2016;49(1):203–12.

31. Petoukhov MV, Franke D, Shkumatov AV, Tria G, Kikhney AG, Gajda M, et al. New developments in the ATSAS program package for small-angle scattering data analysis. J Appl Crystallogr. 2012 Apr 1;45(Pt 2):342–50.

32. Trewhella J, Duff AP, Durand D, Gabel F, Guss JM, Hendrickson WA, et al. 2017 publication guidelines for structural modelling of small-angle scattering data from biomolecules in solution: an update. Acta Crystallogr D Struct Biol. 2017 Sep 1;73(Pt 9):710–28.

33. Franke D, Petoukhov MV, Konarev PV, Panjkovich A, Tuukkanen A, Mertens HDT, et al. ATSAS 2.8: a comprehensive data analysis suite for small-angle scattering from macromolecular solutions. J Appl Crystallogr. 2017 Aug 1;50(Pt 4):1212–25.

34. Svergun DI. Determination of the Regularization Parameter in Indirect-Transform Methods Using Perceptual Criteria. Journal of Applied Crystallography. 1992 Aug 1;25:495–503.

35. Christen B, Perez DR, Hornemann S, Wuthrich K. NMR Structure of the Bank Vole Prion Protein at 20 degrees C Contains a Structured Loop of Residues 165-171. Journal of molecular biology. 2008 Nov 7;383(2):306–12.

36. Lysek DA, Schorn C, Nivon LG, Esteve-Moya V, Christen B, Calzolai L, et al. Prion protein NMR structures of cats, dogs, pigs, and sheep. Proceedings of the National Academy of Sciences of the United States of America. 2005 Jan 18;102(3):640–5.

37. Bernado P, Mylonas E, Petoukhov MV, Blackledge M, Svergun DI. Structural characterization of flexible proteins using small-angle X-ray scattering. J Am Chem Soc. 2007 May 2;129(17):5656–64.

38. Tria G, Mertens HDT, Kachala M, Svergun DI. Advanced ensemble modelling of flexible macromolecules using X-ray solution scattering. Iucrj. 2015 Mar;2:207–17.

39. Proux O, Biquard X, Lahera E, Menthonnex JJ, Prat A, Ulrich O, et al. FAME: a new beamline for x-ray absorption investigations of very-diluted systems of environmental, material and biological interests. Physica Scripta. 2005;2005(T115):970.

40. Filipponi A, Di Cicco A. X-ray-absorption spectroscopy and n-body distribution functions in condensed matter. II. Data analysis and applications. Physical review B, Condensed matter. 1995 Dec 1;52(21):15135–49.

41. Filipponi A, Di Cicco A, Natoli CR. X-ray-absorption spectroscopy and n-body distribution functions in condensed matter. I. Theory. Physical review B, Condensed matter. 1995 Dec 1;52(21):15122–34.

42. D’Angelo P, Bottari E, Festa MR, Nolting HF, Pavel NV. X-ray Absorption Study of Copper(II)-Glycinate Complexes in Aqueous Solution. J Phys Chem B. 1998;102:3114–22.

43. Franke D, Jeffries CM, Svergun DI. Correlation Map, a goodness-of-fit test for one-dimensional X-ray scattering spectra. Nat Methods. 2015 May;12(5):419–22.

44. Carter L, Kim SJ, Schneidman-Duhovny D, Stohr J, Poncet-Montange G, Weiss TM, et al. Prion Protein-Antibody Complexes Characterized by Chromatography-Coupled Small-Angle X-Ray Scattering. Biophys J. 2015 Aug 18;109(4):793–805.

45. Giachin G, Biljan I, Ilc G, Plavec J, Legname G. Probing early misfolding events in prion protein mutants by NMR spectroscopy. Molecules. 2013 Aug 07;18(8):9451–76.

46. Bocharova OV, Breydo L, Salnikov VV, Baskakov IV. Copper(II) inhibits in vitro conversion of prion protein into amyloid fibrils. Biochemistry. 2005 May 10;44(18):6776–87.

47. Orem NR, Geoghegan JC, Deleault NR, Kascsak R, Supattapone S. Copper (II) ions potently inhibit purified PrPres amplification. J Neurochem. 2006 Mar; 96(5):1409–15.

48. McMahon HE, Mange A, Nishida N, Creminon C, Casanova D, Lehmann S. Cleavage of the amino terminus of the prion protein by reactive oxygen species. J Biol Chem. 2001 Jan 19;276(3):2286–91.

49. Yamamoto N, Kuwata K. Difference in redox behaviors between copper-binding octarepeat and nonoctarepeat sites in prion protein. J Biol Inorg Chem. 2009 Nov;14(8):1209–18.

50. Vazquez-Fernandez E, Vos MR, Afanasyev P, Cebey L, Sevillano AM, Vidal E, et al. The Structural Architecture of an Infectious Mammalian Prion Using Electron Cryomicroscopy. PLoS Pathog. 2016 Sep; 12(9):e1005835.

51. Eigenbrod S, Frick P, Bertsch U, Mitteregger-Kretzschmar G, Mielke J, Maringer M, et al. Substitutions of PrP N-terminal histidine residues modulate scrapie disease pathogenesis and incubation time in transgenic mice. PLoS One. 2017;12(12):e0188989.

52. Ashok A, Hegde RS. Selective Processing and Metabolism of Disease-Causing Mutant Prion Proteins. PLOS Pathogens. [doi:10.1371/journal.ppat.1000479]. 2009;5(6):e1000479.

53. Christen B, Perez DR, Hornemann S, Wuthrich K. NMR structure of the bank vole prion protein at 20 degrees C contains a structured loop of residues 165-171. J Mol Biol. 2008 Nov 7;383(2):306–12.

54. Cartoni C, Schinina ME, Maras B, Nonno R, Vaccari G, Di Baria MA, et al. Identification of the pathological prion protein allotypes in scrapie-infected heterozygous bank voles (Clethrionomys glareolus) by high-performance liquid chromatography-mass spectrometry. J Chromatogr A. 2005 Jul 15;1081(1):122–6.

55. Sigurdson CJ, Nilsson KP, Hornemann S, Manco G, Fernandez-Borges N, Schwarz P, et al. A molecular switch controls interspecies prion disease transmission in mice. J Clin Invest. 2010 Jul;120(7):2590–9.

56. Rossetti G, Giachin G, Legname G, Carloni P. Structural facets of disease-linked human prion protein mutants: a molecular dynamic study. Proteins. 2010 Dec;78(16):3270–80.

57. Eghiaian F, Grosclaude J, Lesceu S, Debey P, Doublet B, Treguer E, et al. Insight into the PrPC– – >PrPSc conversion from the structures of antibody-bound ovine prion scrapie-susceptibility variants. Proceedings of the National Academy of Sciences of the United States of America. 2004 Jul 13;101(28):10254–9.

58. Lysek DA, Schorn C, Nivon LG, Esteve-Moya V, Christen B, Calzolai L, et al. Prion protein NMR structures of cats, dogs, pigs, and sheep. Proceedings of the National Academy of Sciences of the United States of America. 2005 Jan 18;102(3):640–5.

59. Spevacek AR, Evans EG, Miller JL, Meyer HC, Pelton JG, Millhauser GL. Zinc drives a tertiary fold in the prion protein with familial disease mutation sites at the interface. Structure (London, England: 1993). 2013 Feb 5;21(2):236–46.

60. Evans EG, Pushie MJ, Markham KA, Lee HW, Millhauser GL. Interaction between Prion Protein’s Copper-Bound Octarepeat Domain and a Charged C-Terminal Pocket Suggests a Mechanism for N-Terminal Regulation. Structure (London, England: 1993). 2016 Jul 6;24(7):1057–67.

61. Sonati T, Reimann RR, Falsig J, Baral PK, O’Connor T, Hornemann S, et al. The toxicity of antiprion antibodies is mediated by the flexible tail of the prion protein. Nature. 2013 Sep 5;501(7465):102–6.

