## Supplementary Table 1, Supplementary Figure 1, 2 and 3 for "Deciphering copper coordination in the animal prion protein amyloidogenic domain"

**Table S1**. SAXS results for the individual proteins investigated in this study

***(a) Sample details.***

| | BvPrP | OvPrP ARR | OvPrP VRQ |
|---|---|---|---|
| Organism | *Myodesglareolus* (Bank vole) | *Ovisaries* (Sheep) | |
| Protein expression system | *Escherichia coli* BL21 DE3 | | |
| UniProt sequence ID (residues in construct) | Q8VHV5 (90-231) | P23907 (94-233) | P23907 (94-233) |
| Genetic polymorphism | wild-type | ARR (A136, R154, R171) | VRQ (V136, R154, Q171) |
| Resistance to TSE | susceptible | resistant | susceptible |
| Extinction coefficient ($M^{-1}$ $cm^{-1}$) | 26025 | 23380 | 23380 |
| Particle contrast from sequence and solvent constituents, $\Delta\rho$ ($\rho_{protein}$ – $\rho_{solvent}$; $10^{10}$; $cm^{-2}$) | 3.061 (12.548 - 9.487) | 3.027 (12.515-9.488) | 3.022 (12.510-9.488) |
| Specific volume from chemical composition (v, $cm^3$ $g^{-1}$) | 0.720 | 0.722 | 0.723 |
| Calculated monomeric $M_r$ from sequence (Da) | 16139.94 | 16084.92 | 16084.91 |
| SEC column | Superdex 75 10/300 GL (GE Healthcare) | Superdex 200 1/150 GL (GE Healthcare) | |
| Injected volume (µL) | 250 | 50 | 50 |
| Loading concentration (mg/mL) | 10 | 7 | |
| Frame range used for data analysis | 1963-2030 (*apo* and copper-loaded) | 682-669 (*apo*), 873-925 (copper-loaded) | 630-670 (*apo*), 865-905 (copper-loaded) |
| Flow rate (mL/min) | 0.5 (*apo* and copper-loaded) | 0.2 (for *apo*) and 0.15 (copper-loaded) | |
| Solvent details | 25 mM NaOAc, 250 mM NaCl, pH 5.5 | | |

***(b) SAXS data collection parameters.***

| | |
|---|---|
| Instrument | ESRF BM29 |
| Wavelength (Å) | 0.99 |
| q-range ($Å^{-1}$) | 0.004 – 0.49 |
| Sample-to-detector distance (m) | 2.864 |
| Exposure time (sec) | 1/frame |
| Temperature (º C) | 20 |
| Detector | Pilatus 1M |
| Flux (photons/s) | $2 \cdot 10^{12}$ |
| Beam size (µ$m^2$) | 700 · 700 |
| Sample configuration | 1.8 mm quartz glass capillary |
| Absolute scaling method | Comparison to water in sample capillary |
| Normalization | To transmitted intensity by beam-stop counter |
| Monitoring for radiation damage | Control of un-subtracted and scaled subtracted data for systematic changes typical for radiation damage |

***(c) Structural parameters.***

| | BvPrP | | OvPrP ARR | | OvPrP VRQ | |
|---|---|---|---|---|---|---|
| Metal bound | *apo* | Cu(II) | *apo* | Cu(II) | *apo* | Cu(II) |
| **Guinier analysis** | | | | | | |
| - I(0) ($cm^{-1}$) | 0.0202 ±2.04E-05 | 0.01788 ±3.19E-05 | 0.01178 ±5.26E-05 | 0.00838 ±1.12E-05 | 0.00964 ±1.76E-05 | 0.009216 ±1.16E-05 |
| - $R_g$ (nm) | 2.42 ±0.052 | 2.38 ±0.073 | 2.31 ±0.01 | 2.25 ±0.06 | 2.31 ±0.090 | 2.27 ±0.091 |
| - $q_{min}R_g$ – $q_{max}R_g$ (nm) | 0.0824-0.3825 (range 55-125) | 0.0818-0.2826 (range 54-106) | 0.0310-0.3145 (range 29-113) | 0.0445-0.2606 (range 38-102) | 0.0475-0.3140 (range 38-113) | 0.0927-0.3140 (range 57-113) |
| - Mass from I(0) (Dalton); ratio to predicted | 16332.58 (1.01) | 15295.12 (0.94) | 15805.80 (0.98) | 15441.28 (0.95) | 15213.62 (0.95) | 15501.24 (0.96) |
| **P(r) analysis** | | | | | | |
| - I(0) ($cm^{-1}$) | 0.0204 | 0.01788 | 0.01192 | 0.008384 | 0.009696 | 0.009248 |

| | | | | | | |
|---|---|---|---|---|---|---|
| - $R_g$ (nm) | 2.41 | 2.37 | 2.32 | 2.26 | 2.308 | 2.24 |
| - $D_{max}$ (nm) | 9.4 | 8.7 | 9.0 | 8.75 | 9.1 | 8.7 |
| - Porod volume ($nm^3$) | 28.54 | 30.13 | 29.44 | 30.75 | 28.74 | 28.77 |
| - $\chi^2$ [total estimate from *GNOM*] | 0.63 | 0.7383 | 0.5417 | 0.6461 | 0.588 | 0.6256 |
| - Mass from I(0) (Dalton); ratio to predicted | 16940.92 (1.05) | 15288.28 (0.94) | 15988.23 (0.99) | 15545.11 (0.96) | 15289.31 (0.95) | 15555.06 (0.96) |
| - Mass estimate [as 0.5 · volume of models (Da)]; (ratio to expected) | 14270 (0.88) | 15065 (0.93) | 14720 (0.91) | 15375 (0.95) | 14370 (0.89) | 14385 (0.89) |

***(d) Software employed for SAXS data reduction, analysis and interpretation.***

| | |
|---|---|
| SAXS data reduction and data processing | EDNA, Primus and Scåtter |
| Extinction coefficient estimate | Protparam |
| Calculation of Δρ and ν values | MULCh 1.1 |
| Shape/bead modelling | DAMMIF *via* ATSAS 2.8.4 |
| Atomic structure modelling | EOM *via* ATSAS on-line (https://www.embl-hamburg.de/biosaxs/atsas-online/) |

***(e)* Shape model-fitting results and atomistic modeling.**

| | **BvPrP** | | **OvPrP ARR** | | **OvPrP VRQ** | |
|---|---|---|---|---|---|---|
| Metal bound | *apo* | Cu(II) | *apo* | Cu(II) | *apo* | Cu(II) |
| ***Ensemble Optimization Method (EOM)*** | default parameters, 10 000 models in initial ensemble, native-like models, constant subtraction allowed | | | | | |
| - Number of rigid domain, PDB model (residues) | 1, 2K56 (170-231) | | 1, 1Y2S (170-231) | | | |
| - $\chi^2$ value | 3.404 | 1.275 | 1.912 | 1.178 | 2.741 | 3.278 |
| - No. of representative structures | 5 | 6 | 5 | 6 | 4 | 5 |
| - $R_{flex}$ ensemble(%) | 83.91 | 82.64 | 82.62 | 80.73 | 77.98 | 79.74 |
| - $R_\sigma$ ensemble | 1.17 | 1.16 | 1.5 | 1.13 | 1.17 | 1.36 |
| - $R_{flex}$ pool (%) | 86.59 | 86.26 | 84.38 | 86.42 | 84.42 | 86.33 |
| - Constant subtracted: | 0.072 | 0.166 | 0.01 | 0.115 | 0 | 0 |
| - Pool, average, $R_g$ (nm) | 2.51 | 2.51 | 2.48 | 2.51 | 2.50 | 2.57 |
| - Pool, average, $D_{max}$ (nm) | 8.55 | 8.41 | 8.34 | 8.42 | 8.47 | 8.69 |
| - Ensemble, >60% of the population, $R_g$ (nm) | 2.22 | 2.07 | 2.14 | 1.93 | 2.21 | 2.15 |
| - Ensemble, >60% of the population, $D_{max}$ (nm) | 8.29 | 6.92 | 7.82 | 6.57 | 8.32 | 7.72 |

***(f)* Small Angle Scattering Biological Data Bank (SASBDB).**

| | **BvPrP** | | **OvPrP ARR** | | **OvPrP VRQ** | |
|---|---|---|---|---|---|---|
| Metal bound | *apo* | Cu(II) | *apo* | Cu(II) | *apo* | Cu(II) |
| SASBDB ID* | SASDEW7 | SASDEX7 | SASDEY7 | SASDEZ7 | SASDE28 | SASDE38 |

* accessible at the following URLs:
https://www.sasbdb.org/data/SASDEW7/l8vjk696cs/
https://www.sasbdb.org/data/SASDEX7/nooyolczz0/
https://www.sasbdb.org/data/SASDEY7/f9s87lr4gi/
https://www.sasbdb.org/data/SASDEZ7/nzhc3dymo7/
https://www.sasbdb.org/data/SASDE28/eymycp0w67/
https://www.sasbdb.org/data/SASDE38/lxoqbwtrrb/

**Figure S1**

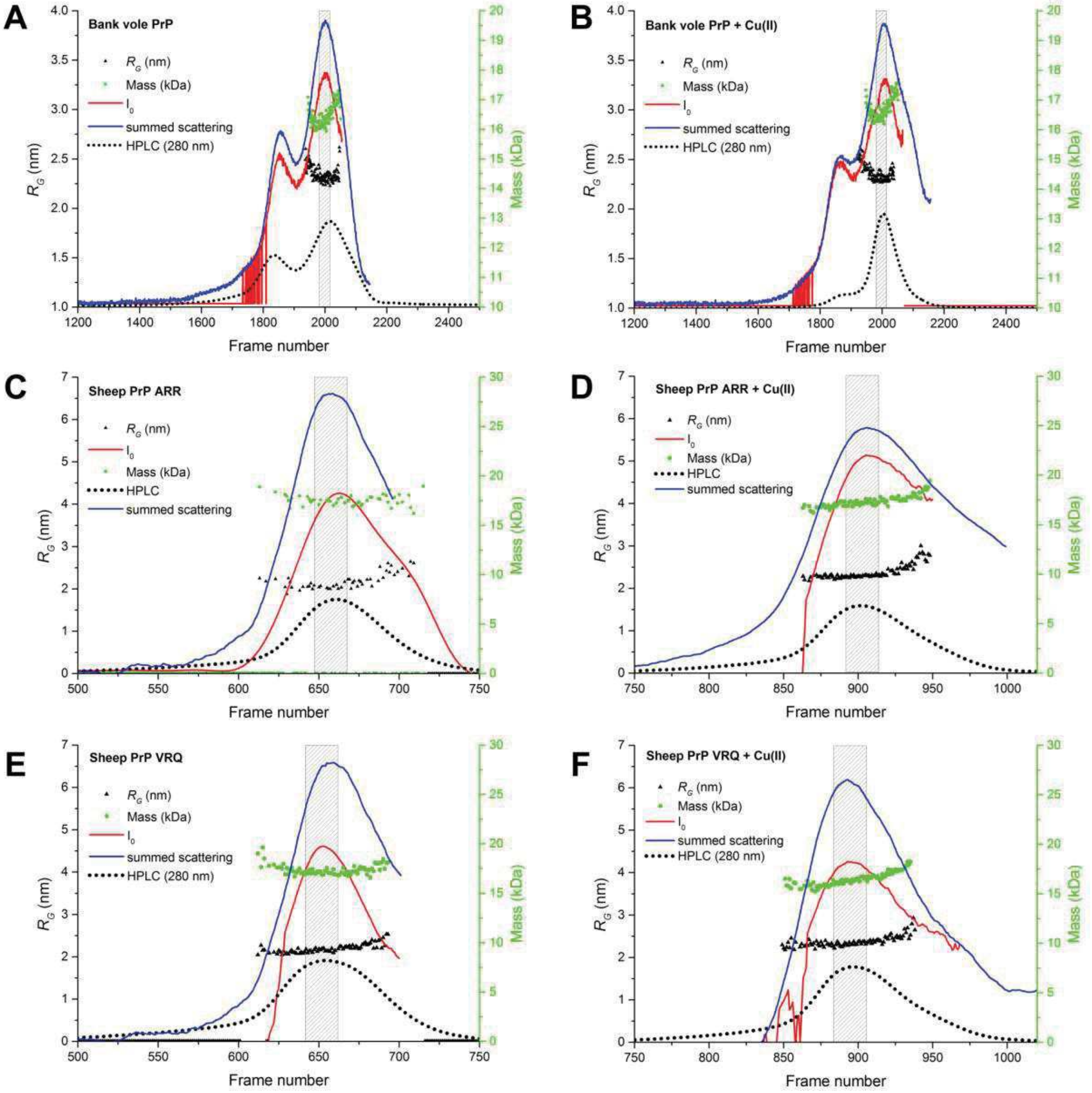


SEC-SAXS chromatograms of Bank vole PrP, sheep PrP ARR and sheep PrP VRQ apo (panels **A**, **C** and **E**, respectively) and Cu(II)-loaded (panels **B**, **D** and **F**, respectively). The blue lines denote the total summed scattering intensity, red lines the forward scattering intensity, I(0), green dots the estimated Mass in kDa based on the correlated volume, black triangles the radius of gyration, $R_g$, and black dots the UV traces at 280 nm from HPLC. The boxes denote regions used for subsequent analysis (see Table S1 for details).

**Figure S2**

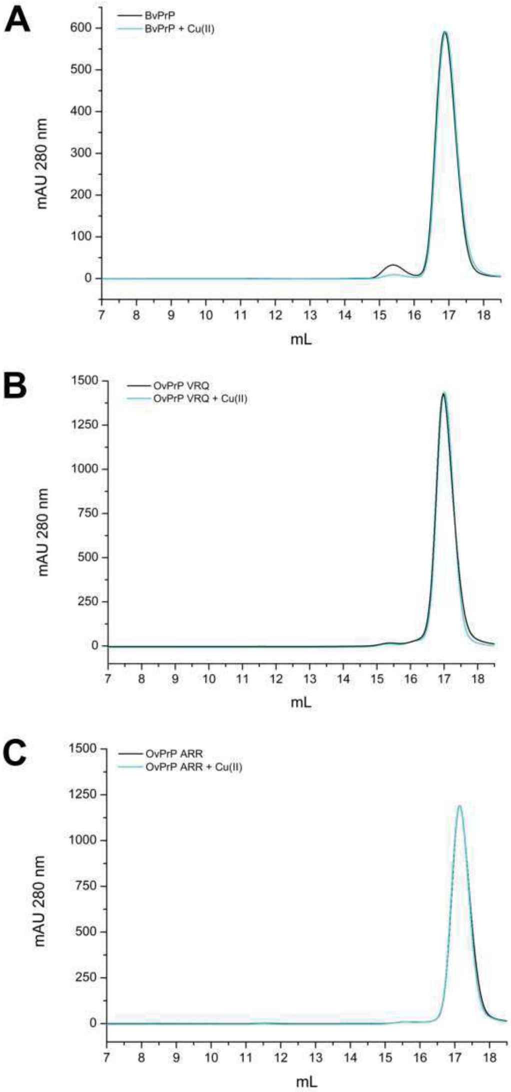


Elution chromatographic profiles of apo and Cu(II)-BvPrP, OvPrP VRQ and OvPrP ARR (panels **A**, **B** and **C**, respectively). All recPrP samples eluted as single, monodispersed peaks at 17.5 mL elution volume (void volume at 7.5 mL, column GE Superdex 200 Increase 10/300).

**Figure S3**

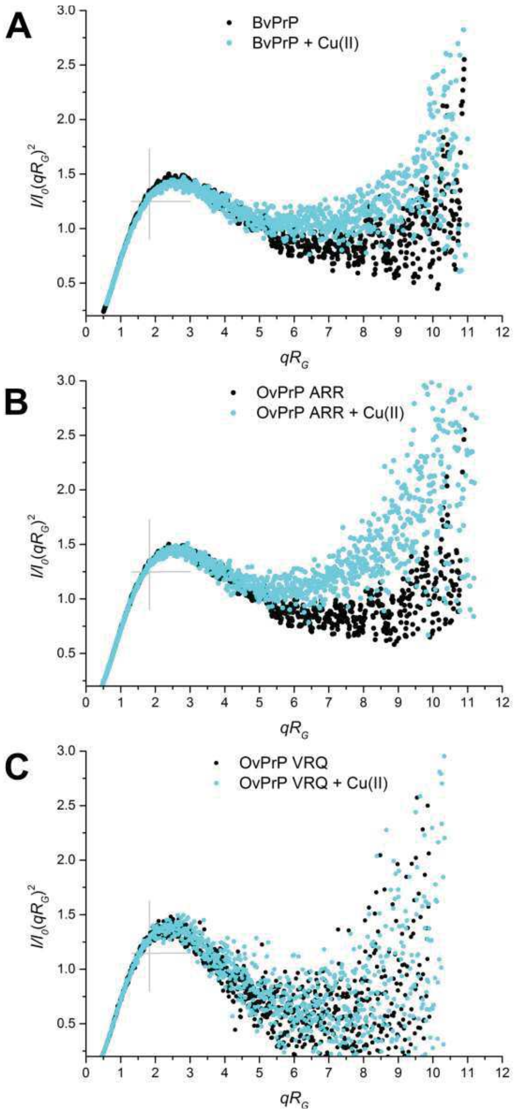


Dimensionless Kratky plots for apo and Cu(II)- BvPrP, OvPrP ARR and OvPrP VRQ, respectively.
